# A novel LC-MS/MS method development and validation for the determination of tezepelumab in rat plasma and its application in rat pharmacokinetic studies

**DOI:** 10.1101/2022.12.14.520442

**Authors:** Madhusudhana Reddy Nimmakayala, Deepti Kolli, J.V. Shanmukha Kumar

## Abstract

By developing a quick, accurate, reproducible, and straightforward LC-MS/MS system and employing trastuzumab as an internal standard (IS), tezepelumab (TZP)’s quantification was achieved. This study explores the most recent advancement in bioanalytical LC-MS/MS technologies utilizing a 150×4.6 mm, 3.5µ Waters Symmetry C18 column that was set for isocratic mode at ambient temperature. Methanol (MeoH): 0.1% Formic acid (FA) in 40:60 v/v at 1.0 ml/min was utilized as the mobile phase. The injection volume and runtime were 10 µl and 5 minutes, respectively. TZP’s retention time (RT) was 1.944 Minutes, and a chromatographic duration total of 5.0 min. With a correlation value (r^2^) of 0.99971, the technique was validated for Tezepelumab throughout a linear range of 6.00–120.00 ng/mL. Findings for precision, accuracy, recovery, matrix effect, and stability all occurred within acceptable limits. Below visual abstract is the concise summary of the article.

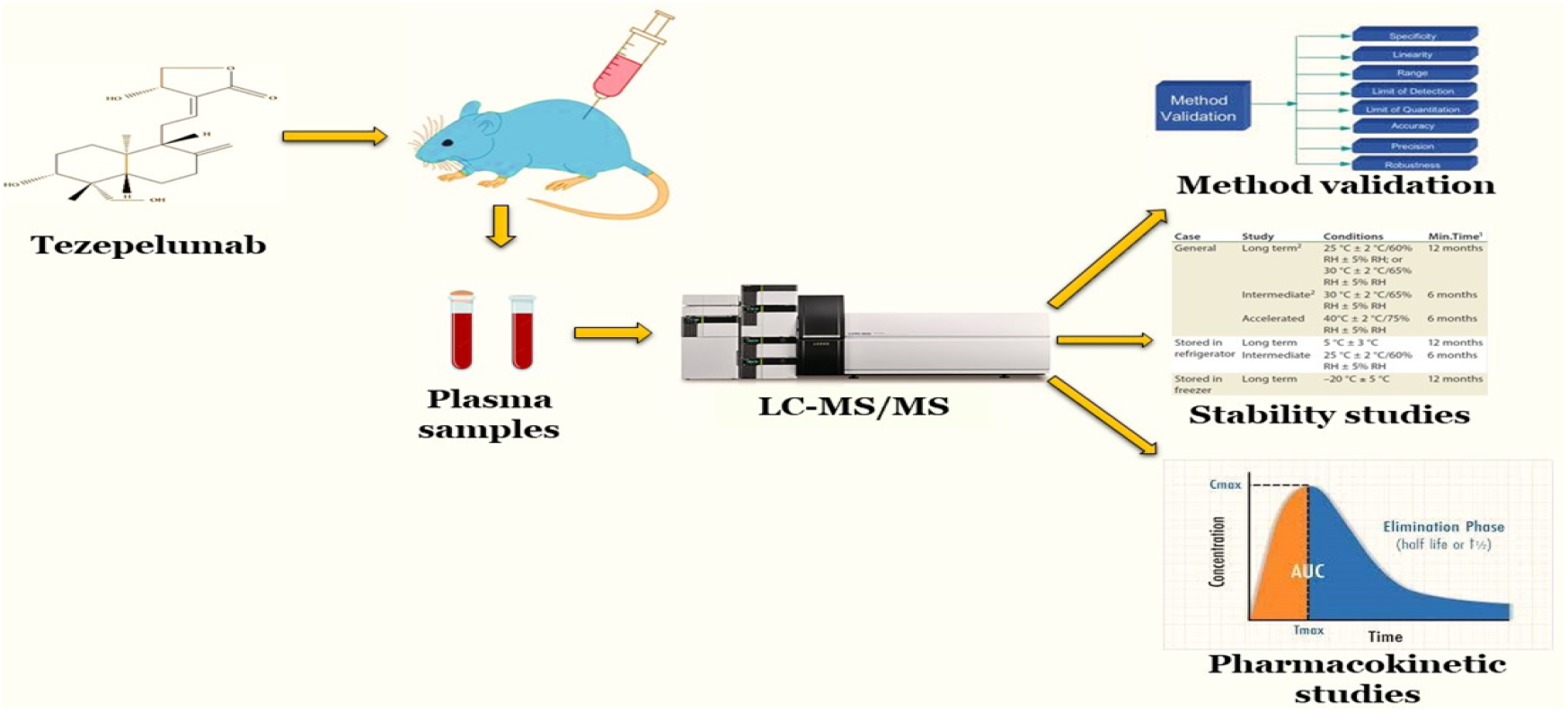

## Introduction

A human monoclonal antibody called tezepelumab (TZP), marketed under the trade name Tezspire, is prescribed for treating asthma.^[1][3][4][6]^ TZP inhibits thymic stromal lymphopoietin,^[1]^ an epithelial mediator implicated in onset as well as worsening of respiratory inflammation.^[8]^ TZP has a chemical formula of C_6400_H_9844_N_1732_O_1992_S_52_, a molecular mass of 144590.40 gmol^-1^. Pharyngitis and arthralgia are among the most frequent adverse reactions. ^[5]^ TZP received medical approval for usage in the US in the year 2021^[1][2]^ and Europe in 2022. ^[3] [7]^ According to a literature review, TZP has not yet been the subject of any research.

## Materials and Methodology

### Chemicals and materials

Samples of TZP and trastuzumab from Biocon in Bangalore. All remaining ingredients, including LCMS grade formic acid (FA), methanol (MeoH), and acetonitrile (ACN), were purchased from Merck Chemical Division in Mumbai. The Milli-Q water purification system’s water was utilized during the whole investigation.

### Instrumentation

For analysis, a Waters, Alliance e-2695 version HPLC equipped with a column oven, auto sampler, and degasser was used. The SCIEX QTRAP 5500 mass spectrometer, which has an electrospray ionization (ESI) interface, was connected to the HPLC system. The results from the chromatogram were interpreted using SCIEX software. The Multiple reaction monitoring (MRM) mode was adopted to record the transformation of protonated precursors to final ions at m/z 147.03 → 82.61 amu and 148.72 → 120.25 amu for sample and IS, correspondingly. The source-dependent variables that were retained for the sample and IS were as follows: GS1: 50.00 psi, GS2: 50.00 psi, IS voltage: 5500.00 V, turbo heater temperature: 550.00°C, collision activation dissociation: 7.00 psi, curtain gas: 20.00 psi. The compound-dependent factors such as decluttering potential were adjusted at 40.00 V, entrance potential, collision energy, and cell exit potential were 10.00 V, 15.00 V, and 7.00 V, accordingly.

### Preparation of standard stock and plasma samples

Required volume of ACN was dissolved to create the standard stock solution of TZP and IS (6 mg/100mL). The diluting solvent (MeOH: 0.1% FA 40:60 v/v) was used to create successive dilution series from stock solutions to spike in plasma in order to generate calibration curve (CC) of standard along with quality control (QC) of sample. Eight non-zero values of 6.00 ng/mL, 15.00 ng/mL, 45.00 ng/mL, 60.00 ng/mL, 75.00 ng/mL, 90.00 ng/mL, and 120.00 ng/mL made up the CC standards.

The QC samples were made up of TZP concentrations of 6.00 ng/mL for the lower limit of quantitation QC (LLOQQC), 30.00 ng/mL for the low-QC (LQC), 60.00 ng/mL for the middle-QC (MQC), and 90.00 ng/mL for the high-QC (HQC). 300 µL of spiked samples were added to polypropylene tubes (that had already been labelled) following bulk spiking. With the exception of 30 samples every one of LQC and HQC that were shifted for preservation in cell frost deep freezer (temp: −17°C to −27°C) in order to generate protracted stability at −22°C ± 5°C, the CC standard as well as QC sample were retained in low temperature freezer (temp: −55°C to −75°C). The technique validation was completed using these samples.

### Extraction procedure

Plasma samples that have been centrifuged and processed should be labeled with time frames. 300 μL of diluent was added to 200 μL of plasma and thoroughly mixed. Estimated 0.5 mL of ACN was combined, followed by extensively vortexing, then swirled for fifteen to twenty minutes at 4000 rev/min to separate all the proteins. The supernatant was filled in a vial before injecting it into the chromatogram.

### Procedure for preparing Mobile phase & Buffer

1 mL of FA and 1000 mL of deionized water were combined together and then the solution was passed through a 0.45 µ filter membrane. 40: 60 v/v mixture of MeOH and 0.1% FA was mixed followed by filtration using 0.45 µ filter paper.

### Preparation of extracted sample

The requisite amount of plasma samples was withdrawn from deep freezer, defrosted at ambient temperature, and the tubes were vortexed. Place the pre-labeled sterile tubes in correct order according to batches; a fraction of 200.000 µL of plasma was blended with 300.000 µL of MeOH, and swirled vigorously for 15 minutes. Following this, 500.000 µL each of diluent, STD stock, and IS stock were blended together, and the solution was mixed thoroughly for 15 min before being vibromaxed for 5 min at 2500 rpm. The solution was centrifuged for five minutes at 4000 rpm and 10°C. Roughly 1.000 ml of supernatant was collected.

### Preparation of un extracted sample

Put 500.000 µl of the STD Stock solution in tubes that have already been labeled. 500.000 µL of the IS working solution should be added, then vortexed to blend properly. Add 1000.000 microliters of mobile phase followed by vortexing. Move the necessary volume into auto-sampler vials that have already been labeled, then inject 10.000 µL into the column as per optimized bio-analytical conditions tabulated in Table 1.

**Table 1.** Optimized Bio-analytical Conditions.

| Parameter | Description |
| --- | --- |
| Column | Water symmetry C<sub>18</sub>, 150mm x 4.6mm, 3.5µm |
| Mobile phase | Methanol : 0.1% Formic acid 40:60 v/v |
| Flow rate | 1.0 ml/min |
| Injection volume | 10µL |
| Retention time | Tezepelumab 1.936 min |
| Run time | 5 min |
| Column oven temperature | Ambient |

### Pharmacokinetic Studies

TZP was extracted from rat plasma using the solvent extraction and partitioning technique. For this, 200 microliters of plasma sample (at the appropriate concentration) were poured into polypropylene tubes and mixed thoroughly. Following the addition of 500 µl of each stock, centrifugation was performed for approximately 10 minutes at 4000 rpm and 20 °C. Every sample’s supernatant was taken in a tube, and it was then vaporized at 40 degrees Celsius for drying. These samples were quickly vortexed after being diluted with 500 µl of diluent and 300 µl of MeOH, and they were subsequently put into auto sampler vials. The Manisha Laboratories animal house facility’s licensing with CPCSEA has been revised for experimental studies on small animals for educational reasons under registration number 1074/PO/Re/S/05/CPCSEA, and this has authorized the pharmacokinetic performance methods.

### Bioanalytical Method validation

The technique was validated for sensitivity, selectivity, linearity, precision, matrix condition, accuracy, reinjection reproducibility, recovery study, and stability.

### Sensitivity & Selectivity

By examining the 6 diverse rat samples and examining interference at respective RTs, sensitivity and selectivity was carried out.

### Matrix effect

To estimate matrix effect, comparability of height area ratios of 6 diverse drug-free samples for TZP was assessed. 6 diverse plasma batches were studied in repeated trials at LQC and HQC concentrations with an adequate precision below 15 %.

### Precision and accuracy

An LLOQQC, LQC, MQC, and HQC level investigation of IS samples was used to assess it. Excluding the LLOQQC, wherein 20% is required, accuracy as well as coefficient of variation (%CV) required to not exceed 15%.

### Recovery

Through the extraction of TZP, the assessment of six samples replicates at every internal control concentration. Then, by correlating the height areas of extracted and non-extracted standards, recovery is investigated.

### Carryover

Carryover refers to the analyte recovered by chromatographic column after reconstitution of this sample using a matrix having a sample concentration upper limit of QC (ULOQC) and beyond.

### Dilution integrity

By injecting the matrix having sample over the ULOQC levels and followed by reconstitution using a blank matrix, the integrity of the dilution must be explicable.

### Stability

By contrasting samples against the new stock sample, the solution stability of the sample was determined. Employing six reruns, the stability in plasma at HQC and LQC levels were ascertained. In accordance with US FDA criteria, the sample was supposed stable if the disparity remained below 15%. The accuracy and stability of spiked plasma maintained at ambient temperature was assessed after being kept in an auto sampler for 24 hours, were evaluated. By studying extract plasma samples that had been pumped promptly opposite samples that had been pumped back following storage in wet extract stability at 2-8°C for 12 h and 18 h, the auto sampler stability (LQC, MQC, & HQC) was assessed. By assessing extract plasma samples that had been pumped right away opposite samples that had been pumped back following storage in dry extract stability at −20 ±3°C for 12 h and 18 h, the reinjection reproducibility was assessed. Freeze-thaw stability was investigated through assessment of freshly spiked IS samples opposite to steadiness samples that had undergone 3 cycles of freezing-thrawing. The concentrations achieved the subsequent day were studied opposite to original concentrations for assessing long-term stability.

High selectivity and recoveries were seen when applying the solvent extraction and partitioning technique. By the application of optimized chromatographic settings, extraction techniques, and detecting factors, TZP in rat plasma may be precisely and accurately detected in less time during the assessment. Figs 1 and 2 depict the parent and production mass spectra of TZP and IS.

**Fig 1.**
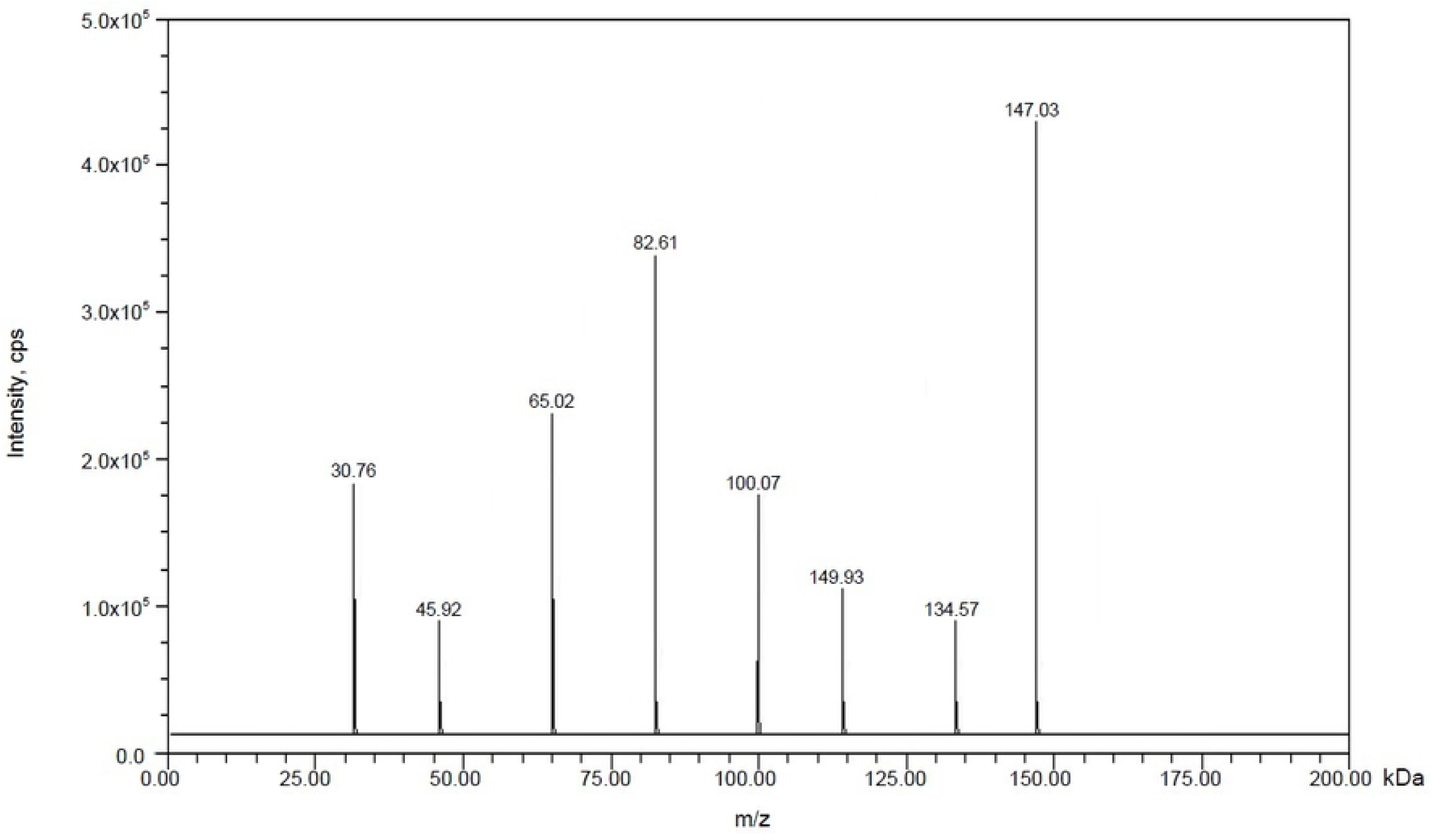
Mass spectrum of fragmentation pattern of Tezepelumab (147.03 & 82.61)

**Fig 2.**
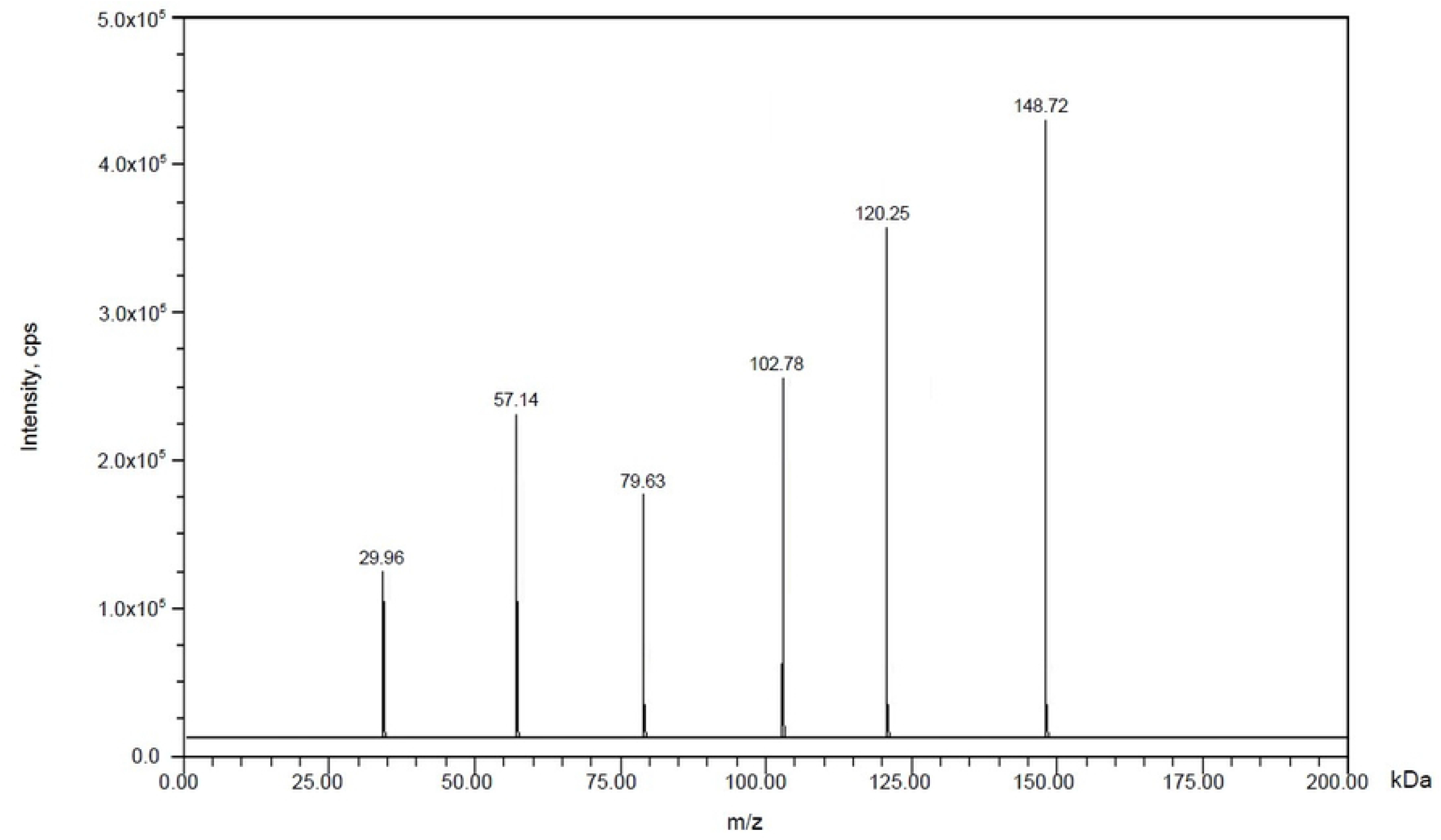
Mass spectrum of fragmentation pattern of IS (148.72 & 120.25)

## Results and Discussion

### Specificity

The chromatographs of blank plasma samples, standard and IS, are depicted in Fig 3, Figs 4 and 5, respectively. There were no interfering peaks visible in the obtained chromatographs.

**Fig 3.**
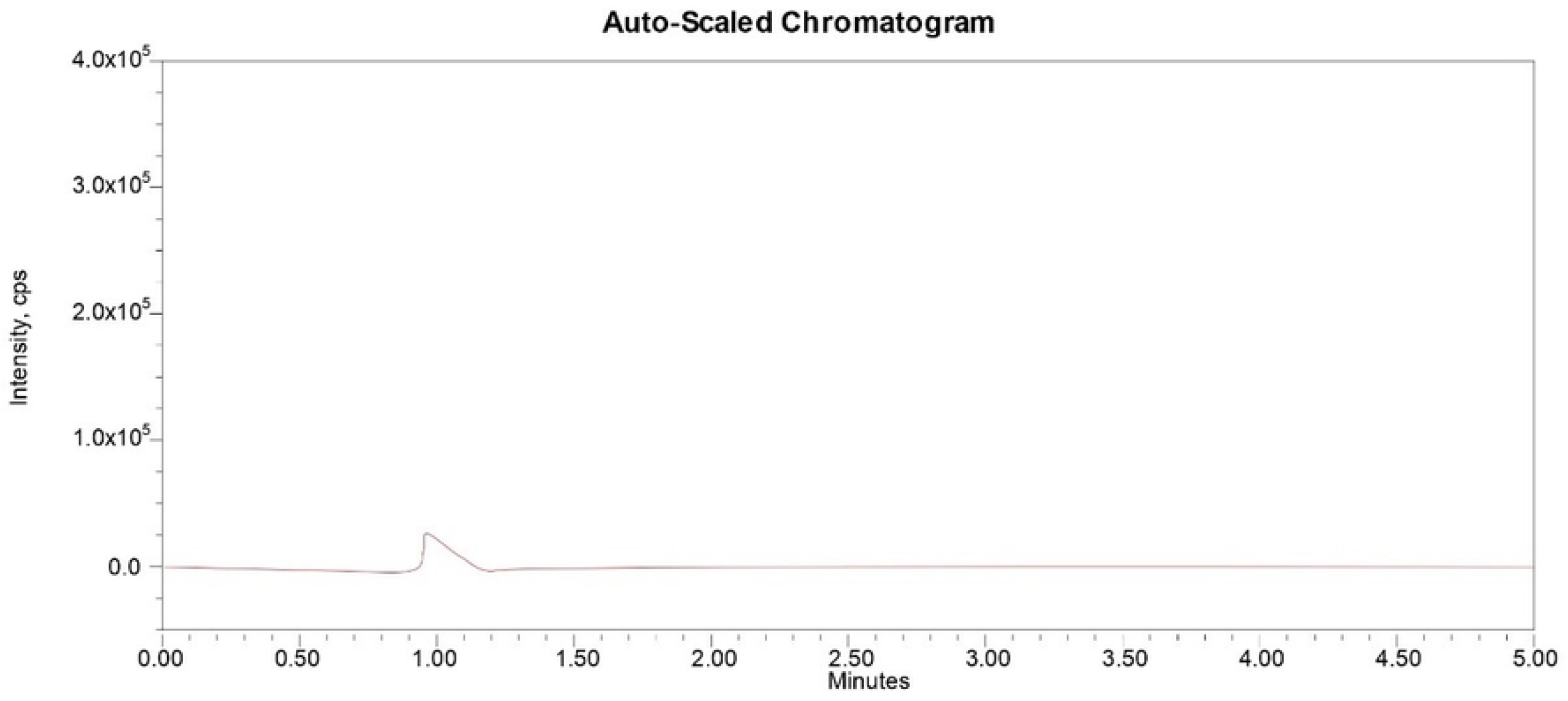
Blank plasma sample chromatogram.

**Fig 4.**
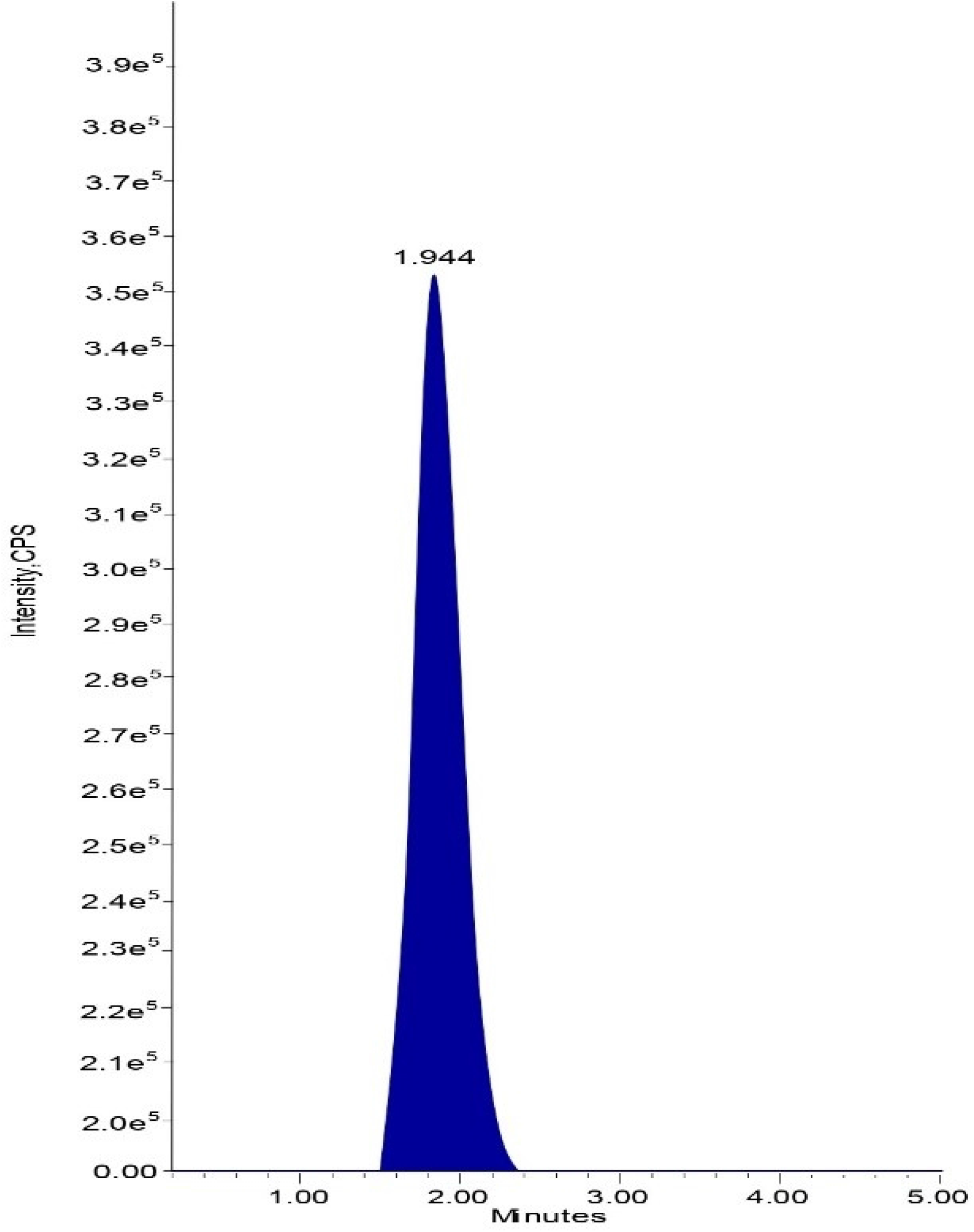
Standard chromatogram of Tezepelumab.

**Fig 5.**
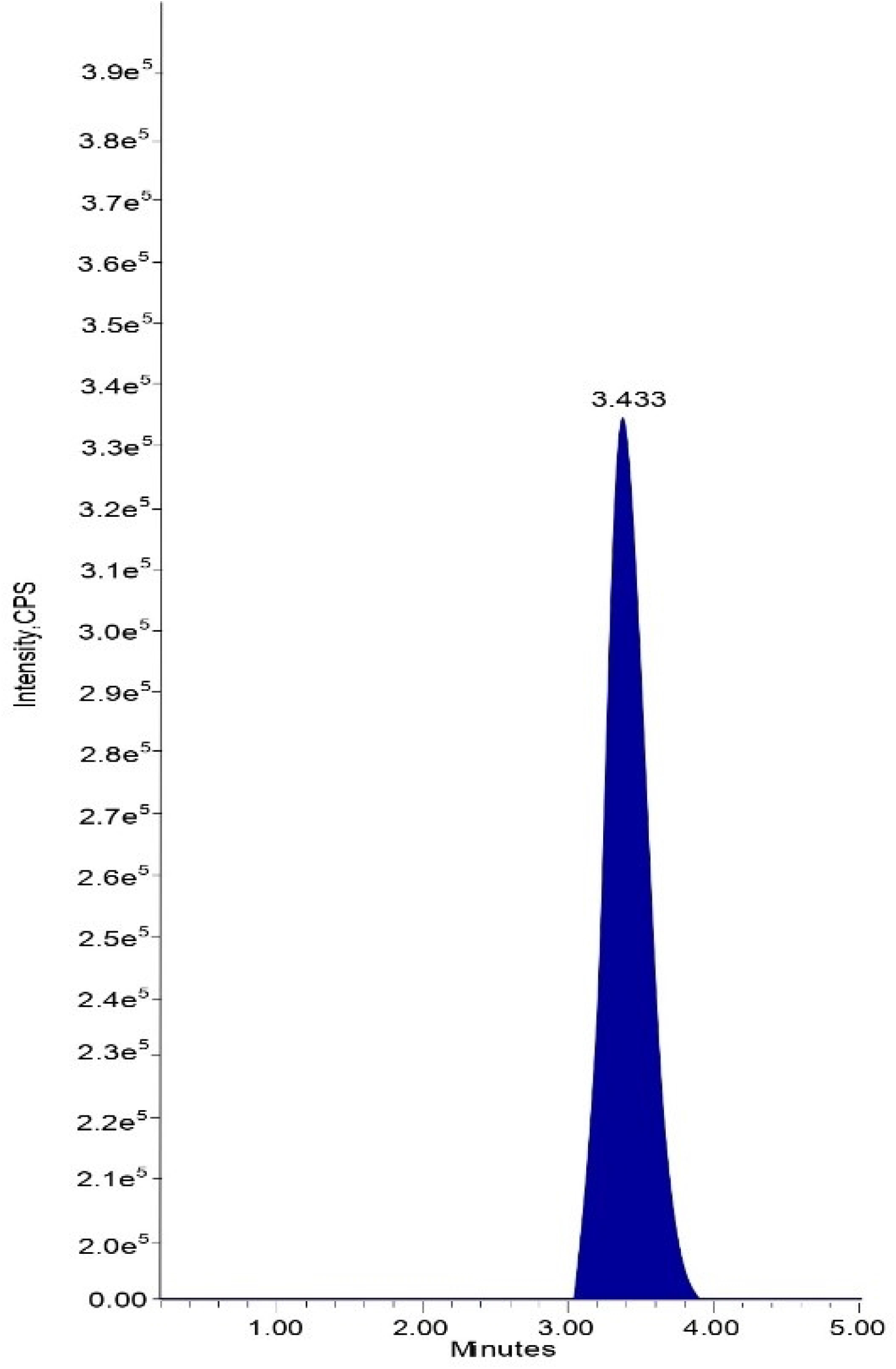
Standard chromatogram of IS.

### Matrix Effect

For TZP LCMS/MS, the % RSD for ionic suppression/upregulation was computed to be 1.0 %, signifying that with these conditions, the matrix effect on sample ionizing falls below an acceptable level. TZP’s LQC and HQC in matrix effect scored 98.46% and 98.55% respectively, and %CV was 1.77 and 0.52, accordingly. It shows that the matrix effect on sample ionizing falls below an appropriate limit of ionization.

### Linearity

At the TZP concentrations spanning between 6-120 ng/mL, the CCs were linear. It was 0.999 for the average correlation coefficient. The proportion of the sample and IS peak areas was used to quantify samples. Plasma concentration was plotted across peak area ratios. Table 2 has the TZP linearity data, and Fig 6 displays their calibration graphs.

**Table 2.** Linearity Results of Tezepelumab.

| **Final conc. in ng/ml** | **Response** | **Area response ratio** |
| --- | --- | --- |
| **0** | 0 | 0.0 |
| **6.00** | 0.356 | 0.105 |
| **15.00** | 0.875 | 0.262 |
| **30.00** | 1.754 | 0.522 |
| **45.00** | 2.537 | 0.755 |
| **60.00** | 3.554 | 1.069 |
| **75.00** | 4.251 | 1.276 |
| **90.00** | 5.253 | 1.558 |
| **120.00** | 7.105 | 2.099 |
| **Slope** | 0.0173 |  |
| **Intercept** | 0.00118 |  |
| **R<sup>2</sup> Value** | 0.99917 |  |

**Fig 6.**
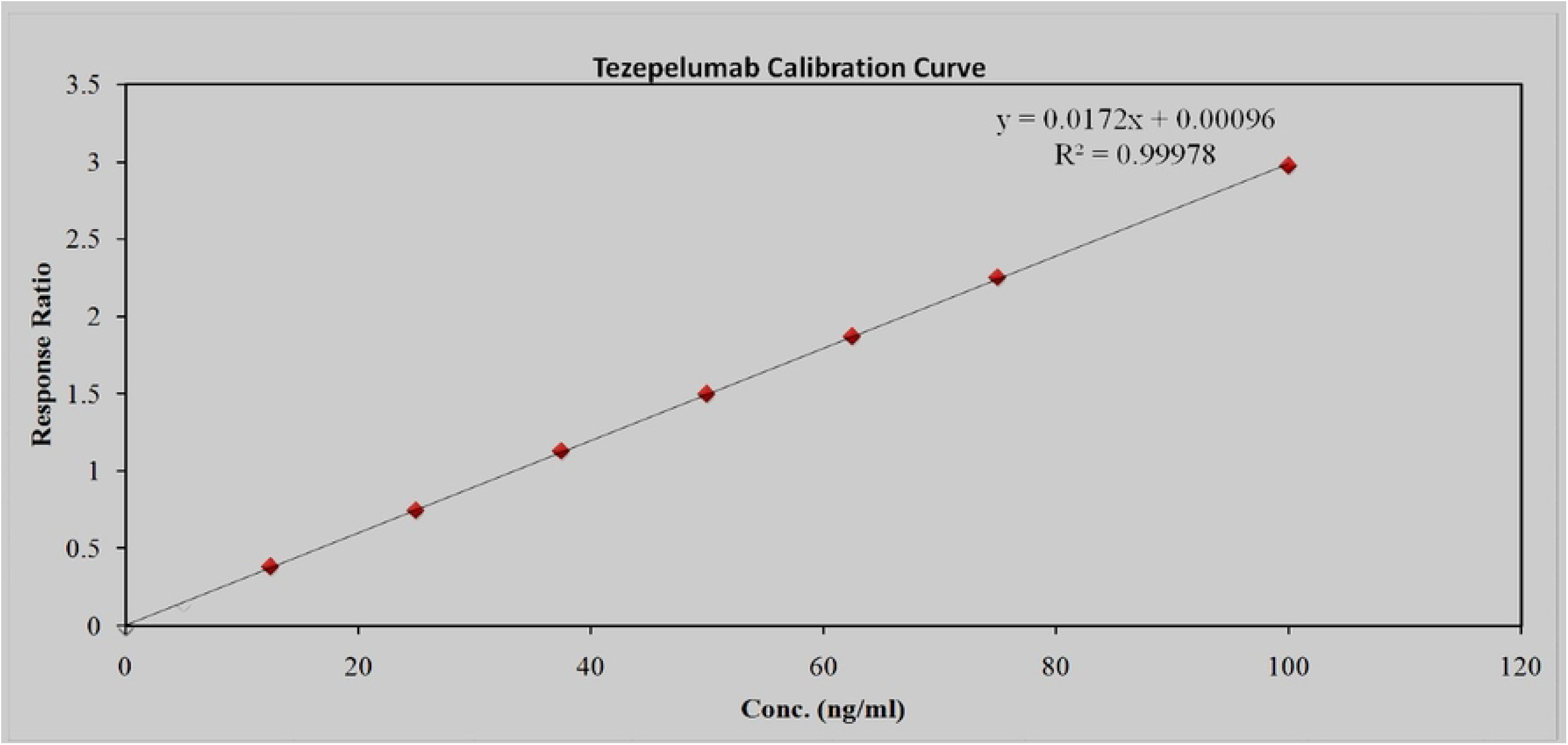
Calibration plot for Tezepelumab.

### Precision & Accuracy

By adding together all of discrete test outcomes from the discrete IS samples, the accuracy and precision were computed. It was clear from the information presented that system was precise and efficient. Table 3 presents TZP precision and accuracy findings.

**Table 3.** Precision and accuracy Results of Tezepelumab.

| Acquisition ID | HQC | MQC | LQC | LLOQQC |
| --- | --- | --- | --- | --- |
|  | Nominal Concentration (ng/ml) |  |  |  |
|  | 90 | 60 | 30 | 6 |
|  | Analyte peak area |  |  |  |
| **1** | 5.280x10<sup>5</sup> | 3.561x10<sup>5</sup> | 1.740x10<sup>5</sup> | 0.369x10<sup>5</sup> |
| **2** | 5.251x10<sup>5</sup> | 3.561x10<sup>5</sup> | 1.796x10<sup>5</sup> | 0.365x10<sup>5</sup> |
| **3** | 5.280x10<sup>5</sup> | 3.587x10<sup>5</sup> | 1.784x10<sup>5</sup> | 0.389x10<sup>5</sup> |
| **4** | 5.282x10<sup>5</sup> | 3.597x10<sup>5</sup> | 1.790x10<sup>5</sup> | 0.367x10<sup>5</sup> |
| **5** | 5.231x10<sup>5</sup> | 3.520x10<sup>5</sup> | 1.778x10<sup>5</sup> | 0.334x10<sup>5</sup> |
| **6** | 5.250x10<sup>5</sup> | 3.523x10<sup>5</sup> | 1.735x10<sup>5</sup> | 0.300x10<sup>5</sup> |
| **n** | 6 | 6 | 6 | 6 |
| **Mean** | 5.262x10<sup>5</sup> | 3.558x10<sup>5</sup> | 1.771x10<sup>5</sup> | 0.354x10<sup>5</sup> |
| **SD** | 0.02132 | 0.03178 | 0.02630 | 0.0318 |
| **% CV** | 0.41 | 0.89 | 1.49 | 8.98 |
| **% Mean Accuracy** | 98.51% | 99.92% | 99.47% | 99.41% |

### Recovery

The approach has high extraction efficiency, as seen by the recovery findings for TZP at LQC, MQC, and HQC concentrations. The %CV for TZP varied between 0.44 and 2.03 and recoveries were 98.16%–105.14% at LQC, MQC, and HQC concentrations. Data implied that proposed approach provides an effectual extraction yield.

### Ruggedness

At LQC, HQC and MQC levels, the % recoveries and CV of TZP as measured by two separate investigators on separate columns met appropriate levels. Data signified that technology is rugged. For TZP, the % recoveries spanned around 98.55% and 100.00%. The %CV figures vary around 0.4 - 1.83. Data implied the ruggedness of the proposed technique.

### Auto sampler carryover

Following sequential loading of LLOQQC as well as ULOQC at the RTs of TZP, the peak area response of TZP was not seen in the blank plasma samples. Consequently, auto sampler carryover is not existent in this approach.

### Stability

For stability investigation, TZP solutions were concocted using solvent and preserved at around 2 and 8 °C. Newly prepared stock solutions and stock solutions concocted a day before were correlated. For 24hrs at 20 °C in the auto sampler and 24h on the benchtop, the plasma stability was steady. Future stability signified that TZP remains stable for a maximum of 24 hrs if maintained at temperature of −30 °C. The Table 4 include TZP outcomes for stability studies.

**Table 4.** Stability studies of Tezepelumab.

| Stability experiment spiked plasma | Concentration | % CV | Mean accuracy |
| --- | --- | --- | --- |
| Benchtop Stability | LOQ-30.0<br>MQC-60.0<br>HQC-90.0 | 1.84 | 98.40 |
|  |  | 0.93 | 100.06 |
|  |  | 0.39 | 97.99 |
| Auto sampler stability |  | 1.43 | 98.46 |
|  |  | 0.87 | 99.75 |
|  |  | 0.61 | 98.29 |
| Freeze-Thaw stability |  | 1.40 | 99.13 |
|  |  | 0.83 | 100.00 |
|  |  | 0.44 | 98.38 |
| Short term stability |  | 0.18 | 95.76 |
|  |  | 0.07 | 96.19 |
|  |  | 0.03 | 97.07 |
| Long term stability Day-28 |  | 0.19 | 86.97 |
|  |  | 0.09 | 87.19 |
|  |  | 0.44 | 89.15 |
| Wet extract 12 hrs. |  | 1.10 | 99.52 |
|  |  | 0.86 | 99.94 |
|  |  | 0.54 | 98.31 |
| Wet extract 18 hr. |  | 1.62 | 98.57 |
|  |  | 0.84 | 99.97 |
|  |  | 0.54 | 98.77 |
| Dry extract 12 hrs. |  | 2.10 | 97.84 |
|  |  | 0.95 | 99.52 |
|  |  | 0.54 | 98.49 |
| Dry extract 18 hrs. |  | 1.68 | 98.06 |
|  |  | 0.73 | 100.34 |
|  |  | 0.42 | 98.29 |

### Pharmacokinetic studies

Six distinct rats were given injections of TZP samples at various intervals, including 2, 4, 6, 8, 12 and 24 days. Samples are then concocted in accordance with the test methodology, loaded into the chromatographic device, and the findings are noted & tabulated in Table 5.

**Table 5.**
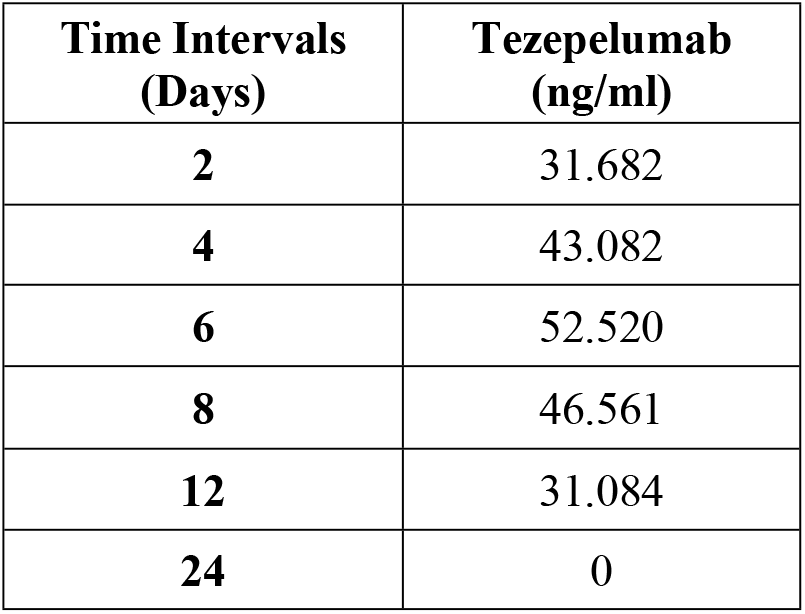
Pharmacokinetic studies of Tezepelumab.

| **Time Intervals**<br>**(Days)** | **Tezepelumab**<br>**(ng/ml)** |
| --- | --- |
| **2** | 31.682 |
| **4** | 43.082 |
| **6** | 52.520 |
| **8** | 46.561 |
| **12** | 31.084 |
| **24** | 0 |

Animals received a single 210 mg dose of TZP injection; samples were drawn 2, 4, 6, 8, 12 and 24 days later. K2 EDTA vacuum blood collection tubes were utilized for acquiring a portion of 5 ml blood at every interval. A pre-injection sample was withdrawn to assess for any noticeable interferences. The plasma was extracted from the obtained samples using centrifugation technique, and it was then kept at −70°C. At diverse concentrations, samples were spiked with IS and assessed along with QC samples. The WinNonlin (Version 5.2) software program was implemented to determine the pharmacokinetic profile of TZP. By incurred sample reanalysis (ISR), the stability of the research samples was determined. Close to C_max_ as well as the elimination phases in the pharmacokinetics, two samples from every subject were chosen for ISR. The percentage difference shouldn’t be beyond 20%, and the samples were thought to be stable. Fig 7 represents recovery graph for Tezepelumab in rat plasma.

**Fig 7.**
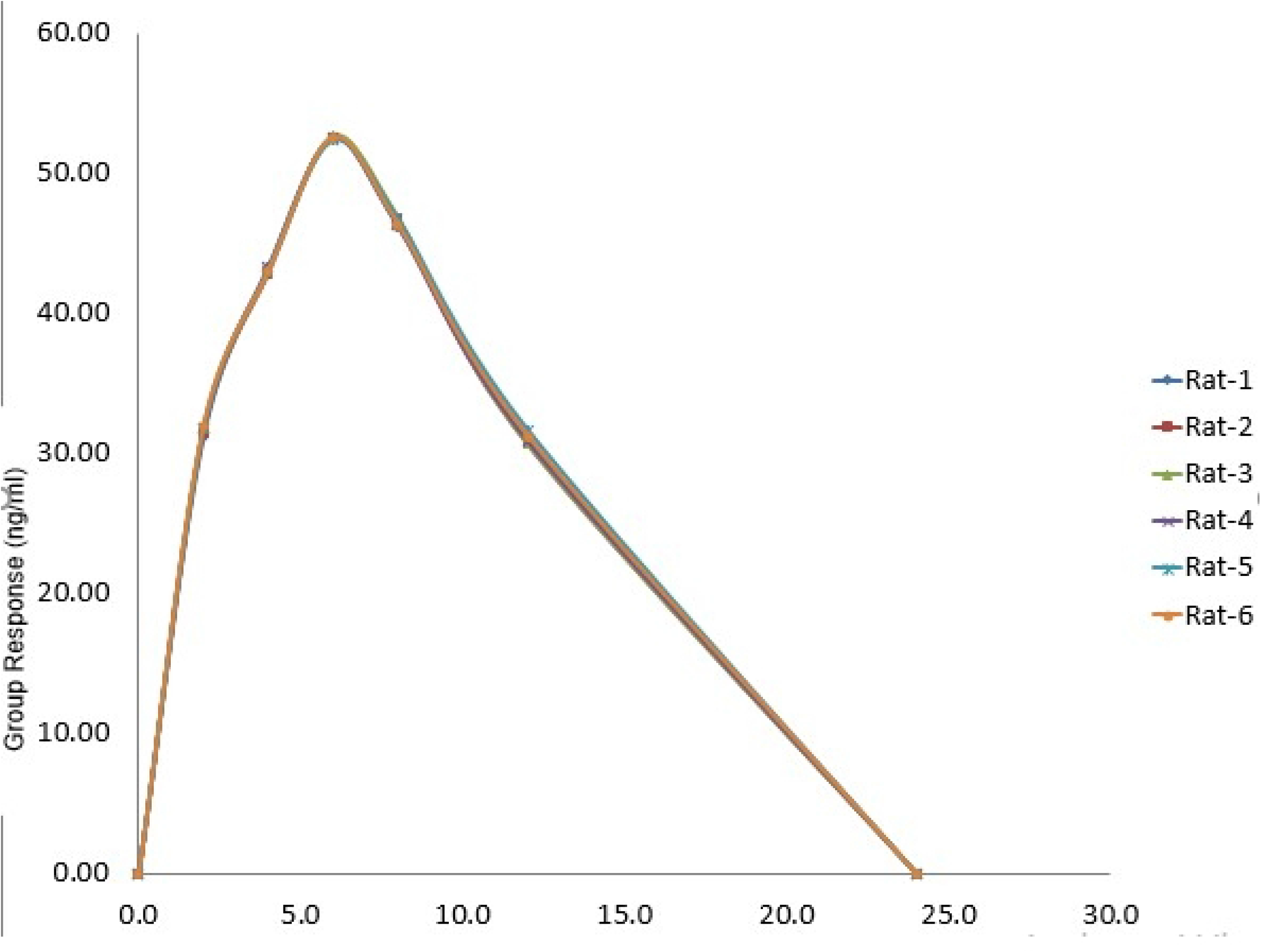
Recovery graph for Tezepelumab in plasma.

## Summary and Conclusion

The goal of this research was to establish an easy, affordable, reliable, and efficient technique for measuring Tezepelumab in LCMS by utilizing Trastuzumab as an internal reference. TZP’s RT was 1.944 minutes, whereas that of IS was 3.433 minutes, for an overall chromatographic duration of 5.0 minutes. Having r^2^ value of 0.999 correlation, the technique is validated for TZP throughout a flexible linear range of 6-120 ng/mL. The accuracy (%CV) for 5 levels was below 15.00 both within and across batches. In accordance with USFDA regulations, this may be verified.

